# Potential therapy for progressive vision loss due to *PCDH15*-associated Usher Syndrome developed in an orthologous Usher mouse

**DOI:** 10.1101/2021.06.08.447565

**Authors:** Saumil Sethna, Wadih M. Zein, Sehar Riaz, Arnaud P. J. Giese, Julie M. Schultz, Todd Duncan, Robert B. Hufnagel, Carmen C. Brewer, Andrew J. Griffith, T. Michael Redmond, Saima Riazuddin, Thomas B. Friedman, Zubair M. Ahmed

## Abstract

Usher syndrome type I (USH1) is characterized by congenital deafness, vestibular areflexia, and progressive retinal degeneration with age. The protein-truncating p.Arg245* founder variant of *PCDH15* has an ~2% carrier frequency among Ashkenazi Jews, accounting for nearly 60% of their USH1 cases. Here, longitudinal ocular phenotyping in thirteen USH1F individuals harboring the p.Arg245* variant revealed progressive retinal degeneration, leading to severe loss of vision with macular atrophy by the sixth decade. Half of the affected individuals met either the visual acuity or visual field loss definition for legal blindness by the middle of their fifth decade of life. Mice homozygous for p.Arg250* (*Pcdh15^R250X^*; equivalent to human p.Arg245*) also have early visual deficits evaluated using electroretinography. Light-dependent translocation of phototransduction cascade proteins, arrestin and transducin, was found to be impaired in *Pcdh15^R250X^* mice. Retinal pigment epithelium-(RPE) specific visual retinoid cycle proteins, RPE65 which converts all-*trans* retinoids to 11-*cis* retinoids and CRALBP that transports retinoids, and key retinoid levels were also reduced in *Pcdh15^R250X^* mice, suggesting a dual role for protocadherin-15 in photoreceptors and RPE. Administration of exogenous 9-*cis* retinal, an analog of the naturally occurring 11-*cis* retinal, improved ERG amplitudes in these mutant mice, suggesting a basis for a clinical trial of exogenous FDA approved retinoids to preserve vision in USH1F patients.

**Summary:** In a preclinical setting studying exogenous retinoids using a novel Usher syndrome mouse model, we describe a potential therapy to treat *PCDH15*-mediated visual dysfunction.

## Introduction

Loss of vision in individuals with USH1, an autosomal recessive disorder, begins towards the end of their first decade of life due to retinitis pigmentosa (RP), eventually leading to near total blindness. Night blindness is an early sign in USH1 subjects followed by constriction of the visual field (tunnel vision) and finally clinical blindness (Vernon, 1969). Characteristic fundus features include pigmentary retinopathy, narrowing of the retinal vessels, and a pale appearance of the optic disk (Toms, Pagarkar et al., 2020). Vestibular dysfunction in USH1 manifests as a delay in development of independent ambulation while hearing loss is usually severe to profound, congenital and sensorineural (Ahmed, Riazuddin et al., 2003b, Smith, Berlin et al., 1994). Cochlear implants can restore auditory perception in USH1 patients (Brownstein, Ben-Yosef et al., 2004a, Pennings, Damen et al., 2006), but presently there is no effective treatment for the vision loss due to RP. Moreover, there is a lack of longitudinal data for the natural history of ocular abnormalities associated with variants of *PCDH15* in humans. Only anecdotal clinical data has been reported thus far (Ahmed, Riazuddin et al., 2001a, Ben-Yosef, Ness et al., 2003, Brownstein, Ben-Yosef et al., 2004b, Jacobson, Cideciyan et al., 2008). Here, we describe the natural history of retinopathy (followed for up to 30 years) in 13 individuals with an Ashkenazi Jewish recessive founder variant of *PCDH15.* Eleven patients from nine families were homozygous for the p.Arg245*, leading to truncation of the encoded protein, protocadherin-15. Two additional patients had compound heterozygous genotypes that included one p.Arg245* allele.

Protocadherin-15 is a member of a large cadherin superfamily of calcium-dependent cell– cell adhesion molecules (Ahmed, Goodyear et al., 2006a, Ahmed, Riazuddin et al., 2001b, van Roy, 2014). Within the vertebrate inner ear, protocadherin-15 is required for the structural maintenance and the mechanotransduction function of the sensory hair cells (Ahmed, Goodyear et al., 2006b, Kazmierczak, Sakaguchi et al., 2007). In the retina, protocadherin-15 is localized to the outer limiting membrane of photoreceptors (PR) and in Müller glia (Reiners, van Wijk et al., 2005, van Wijk, van der Zwaag et al., 2006). We previously reported a reduction of ERG a- and b-waves amplitudes (~40%) at 5 weeks of age in at least two *Pcdh15* alleles in mice (*Pcdh15^av-5J^ and Pcdh15^av-jfb^*) (Haywood-Watson, Ahmed et al., 2006b). However, the exact molecular function of protocadherin-15 in the retina remains elusive. Here, we describe the pathophysiology and function of protocadherin-15 in the retina of a novel murine model. Finally, electroretinogram (ERG) data show significant improvement after treatment of our USH mouse model with 9-*cis* retinal, raising the possibility that exogenous retinoids could preserve vision in *USH1F* patients.

## Results and Discussion

### Spectrum and longitudinal ocular phenotypic data revealed early onset rod-cone dystrophy in USH1 subjects

We reviewed the medical records of 13 patients enrolled in an Institutional Review Board-approved protocol to study Usher syndrome. Subsequent to congenital profound deafness, the first reported symptom was difficulty with vision at night. Ophthalmic manifestations depended on the age of the patient and the stage of retinal degeneration (**Table 1**). Electroretinography recordings were at noise-level for both scotopic and photopic responses, suggesting dysfunctional photoreceptors (**Table 1**). In young patients with early stages of the retinal degeneration, findings included mottling of the retinal pigment epithelium with early pigment redistribution, and mild to moderate vascular attenuation with typically preserved macular reflexes (**Fig. 1a**). As RP progressed, more extensive pigment abnormalities were observed with deposition of bone spicules, severe attenuation of the retinal vasculature, macular atrophic changes, and a waxy pallor of the optic nerve head (**Fig. 1b**). In advanced stages these changes became more prominent and widely distributed throughout the fundus (**Fig. 1c**). Cataracts were common, especially posterior subcapsular opacities. The panels in **Fig. 1d** show the progression of macular atrophic changes over a 12-year period in a patient with compound heterozygous p.Arg245*/p.Arg929* variants (both alleles of *PCDH15* are predicted to cause truncation of the protocadherin-15 protein). Kinetic visual field testing (**Table 1**) showed early loss of the ability to detect the smaller and dimmer target (I1e) in all but one patient (LMG268 #1722 at age 37 years). Early midperipheral scotomas and severe constriction were noted while testing the I4e isopter (target is equal in size to I1e but brighter). Progressive constriction of the V4e visual field isopter (largest and brightest target) is seen in **Fig 1e** where the horizontal diameter is binned by the decade of life. **Fig 1f** shows an increase in logMAR visual acuity, corresponding to a decline in Snellen best-corrected visual acuity (BCVA), starting at the fourth decade of life. Kaplan-Meier survival curves (**Fig 1g**) with parameters corresponding to legal blindness (acuity at 20/200 and visual field limited to 20 degrees) demonstrate severe visual function loss by the fifth decade.

**Table 1:**
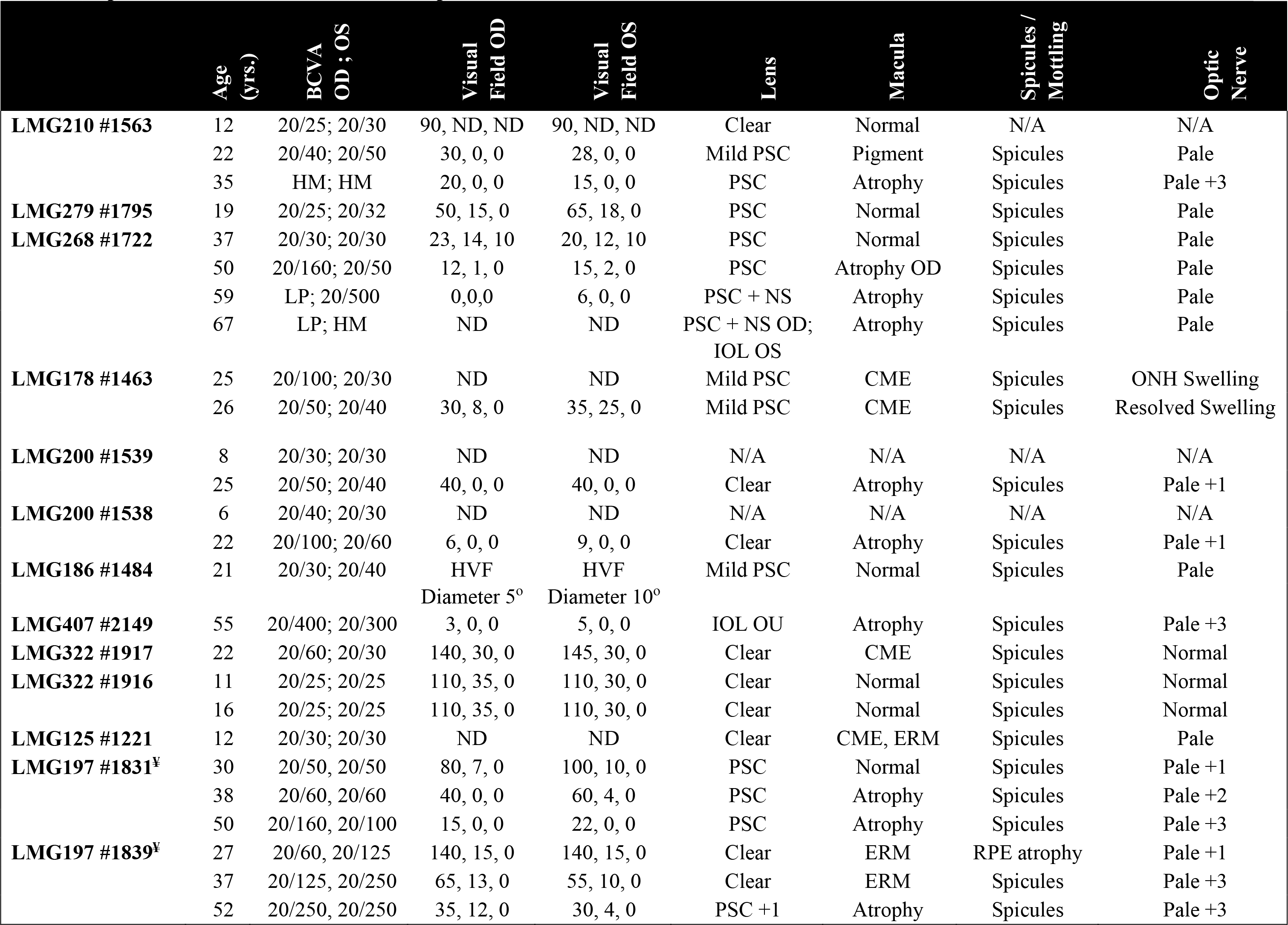
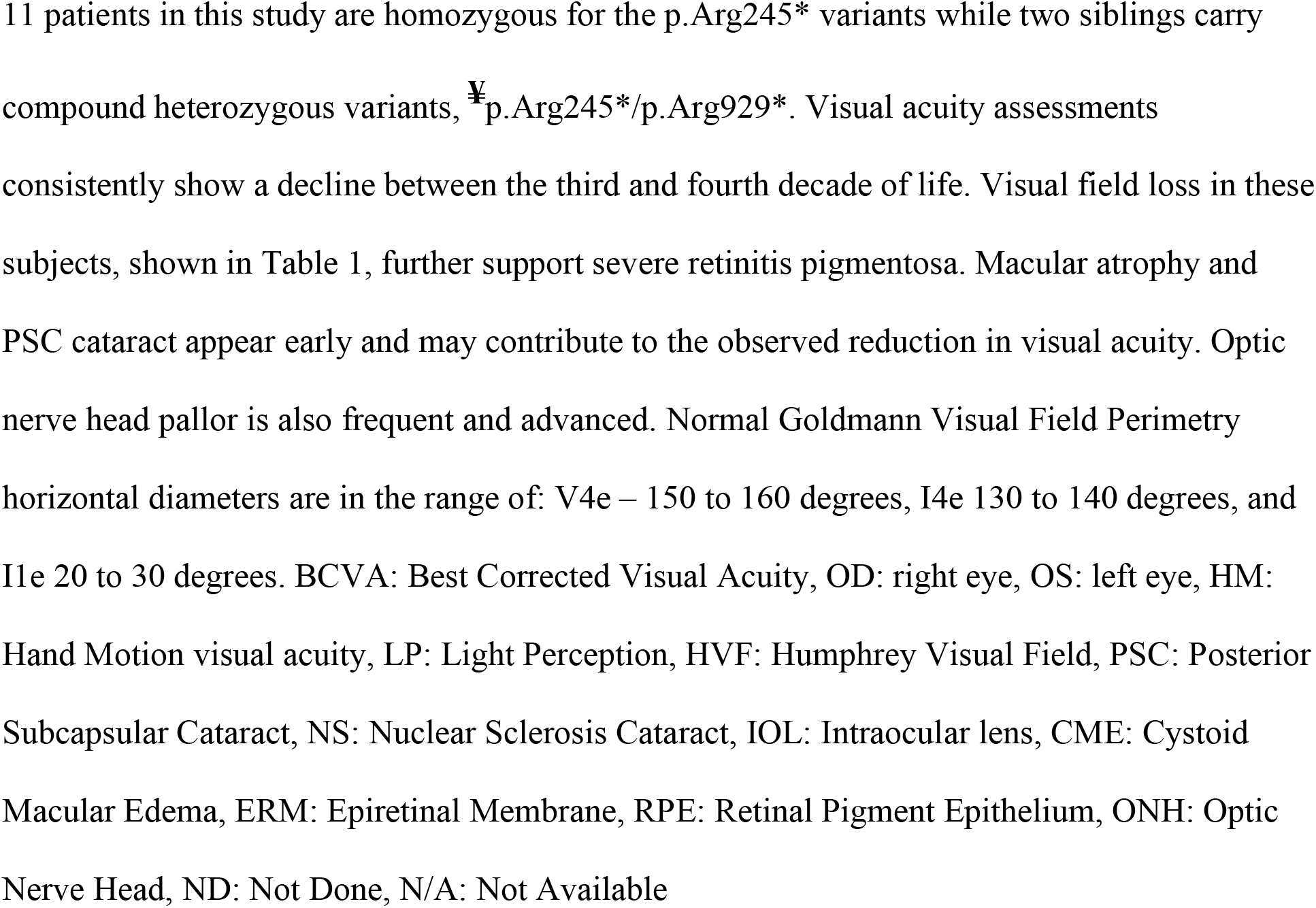
Ophthalmic clinical manifestations of patients with biallelic *PCDH15* mutations.

**Figure 1:**
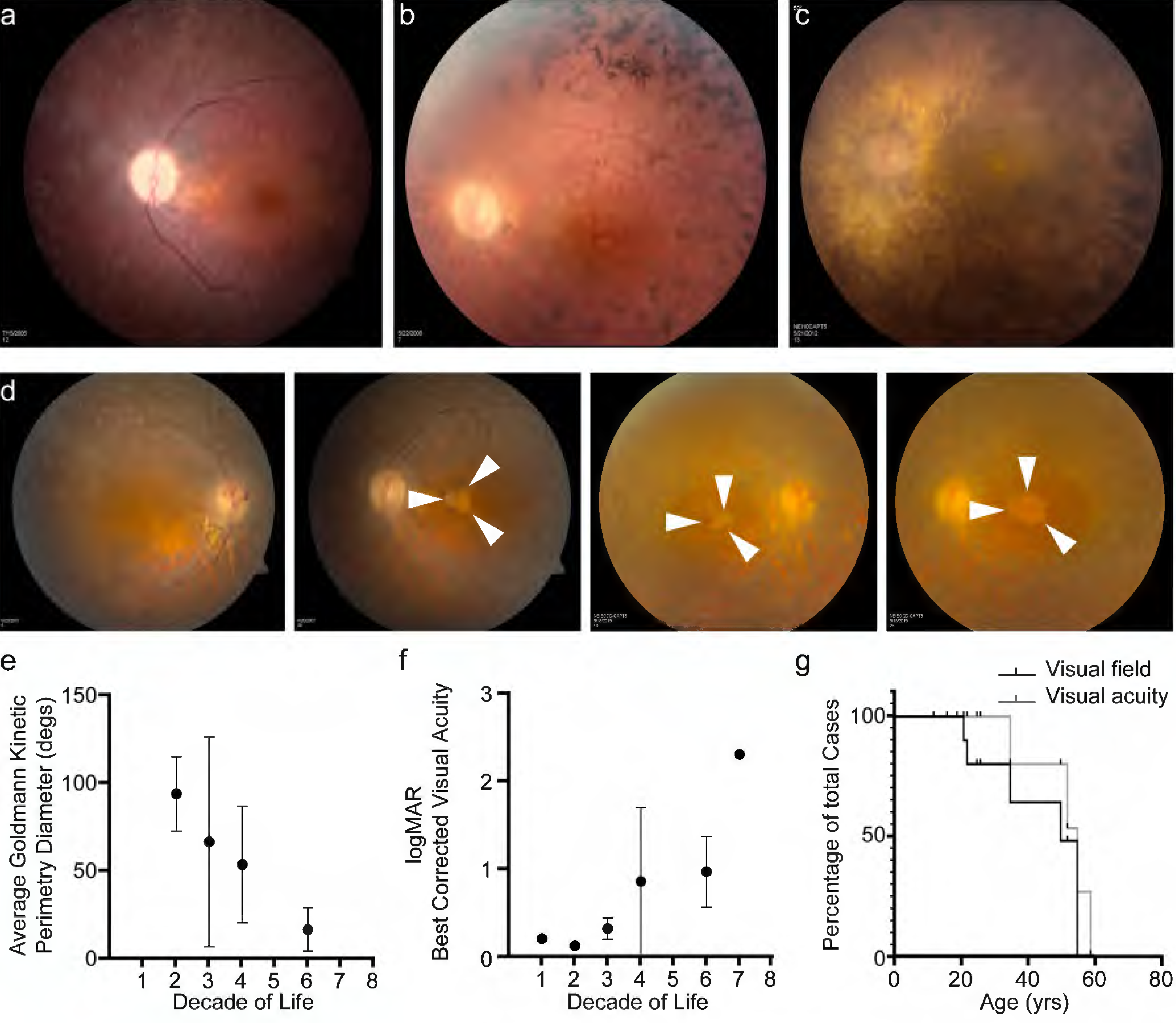
USH1F p.Arg245* spectrum and longitudinal eye phenotype. **a-c** Fundus images depicting the spectrum of retinal findings in p.Arg245* USH1F patients, show mottling of pigment epithelium, attenuation of retinal vasculature and pallor of optic nerve head seen in all three fundus photos. Peripheral bony spicules and macular atrophy are noted **b**. Diffuse atrophy and advanced retinal degeneration are seen in **c**. **d** Longitudinal progression of macular atrophic changes (arrowheads point at edge of macular atrophic area) over a 12-year period in a USH1F patient who is compound heterozygous for p.Arg245*/p.Arg929*. **e** Mean and SEM of Goldmann visual field diameters for patients with data binned by decade of life. **f** Mean and SEM of best corrected visual acuity binned by decade of life for all patient visits. **g** Survival analysis curves for visual acuity (logMAR visual acuity >1 i.e. acuity worse than 20/200) and visual field (visual field < 20 degrees in better eye). These values were chosen since they usually denote visual function at legal blindness levels. SEM, standard error of the mean.

### *Pcdh15^R250X^* knockin mutants recapitulates human p.Arg245* Usher phenotype

In order to investigate the precise role of protocadherin-15 in light transduction and the mechanism of visual deficits observed in patients afflicted with the Arg245* pathogenic variant of *USH1F*, we used CRISPR/Cas9 technology to engineer a mouse model with the *Pcdh15^R250X^* variant (**Methods**) (Cong, Ran et al., 2013). The p.Arg250* variant is located in an exon common to all *Pcdh15* transcripts and, consequently, is predicted to cause a complete loss of all known protocadherin-15 isoforms (Ahmed et al., 2006b). Indeed, with immunostaining of cochlear tissue (**Fig. S1a**) or retinal tissue (**Fig. S3a**), we could not detect protocadherin-15 expression in these tissues from *Pcdh15^R250X/R250X^* mutant mice.

Consistent with previously published *Pcdh15* mouse models (Alagramam, Goodyear et al., 2011, Haywood-Watson, Ahmed et al., 2006a, Senften, Schwander et al., 2006) we detected no auditory-evoked brainstem responses (ABRs) in mutant *Pcdh15^R250X^* mice at P16, the earliest postnatal day that ABRs can be reliably detected (**Fig. S2a**), indicating that they were profoundly deaf. Furthermore, *Pcdh15^R250X^* mutant mice displayed abnormal motor vestibular behaviors such as circling, hyperactivity, and head bobbing. Behavioral tests including exploratory-behavior and tail-hanging tests confirmed that these deaf mice also have a significant vestibular dysfunction (**Fig. S2b, c**). Lastly the *Pcdh15^R250X^* mutant cochlear and vestibular hair cells also had no functional mechanotransduction (**Fig. S2d**), accounting for deafness, and at P60 also showed degeneration of hair cells in the organ of Corti (**Fig. S1b-c**). Taken together, our data indicate that the *Pcdh15^R250X^* mutants recapitulates human p.Arg245* deafness and peripheral vestibular areflexia.

To parallel the visual examinations performed in patients, we assessed the visual function of *Pcdh15^R250X^* mutant mice using full-field electroretinography (ERG). Dark-adapted (scotopic) ERG waves, which are preferentially driven by rod photoreceptors at low light intensity and by rod and cone photoreceptors at high light intensity, showed normal wave architecture albeit with reduced amplitudes (**Fig. S3b**). Quantification showed significant reduction in amplitudes of the a-wave (derived primarily from the photoreceptor layer) and the b-wave (derived from Müller glia and outer nuclear bipolar neurons) in one-month-old *Pcdh15^R250X^* mutant mice as compared to littermate control mice (**Fig. 2a**). Similarly, photopic ERG amplitudes, primarily representing cone-mediated function, were also reduced in one-month-old *Pcdh15^R250X^* mutant mice (**Fig. 2b**). The b- to a-wave ratio (b/a) was similar across genotypes indicating that deficits were manifested mainly at the photoreceptor level (**Fig. S3c**). We then performed ERGs at 2-3 and 6-7 months of age. *Pcdh15^R250X^* mutant mice consistently had lower scotopic and photopic ERG amplitudes compared to controls (**Fig. 2c-f**), indicating that the functional deficits observed at one month were not due to delayed development. To correlate functional deficits with structural integrity of the retina, we performed non-invasive *in vivo* retinal imaging using optical coherence tomography (OCT), which showed no gross structural abnormality in *Pcdh15^R250X^* mutant mice (**Fig. S3d**).

**Figure 2:**
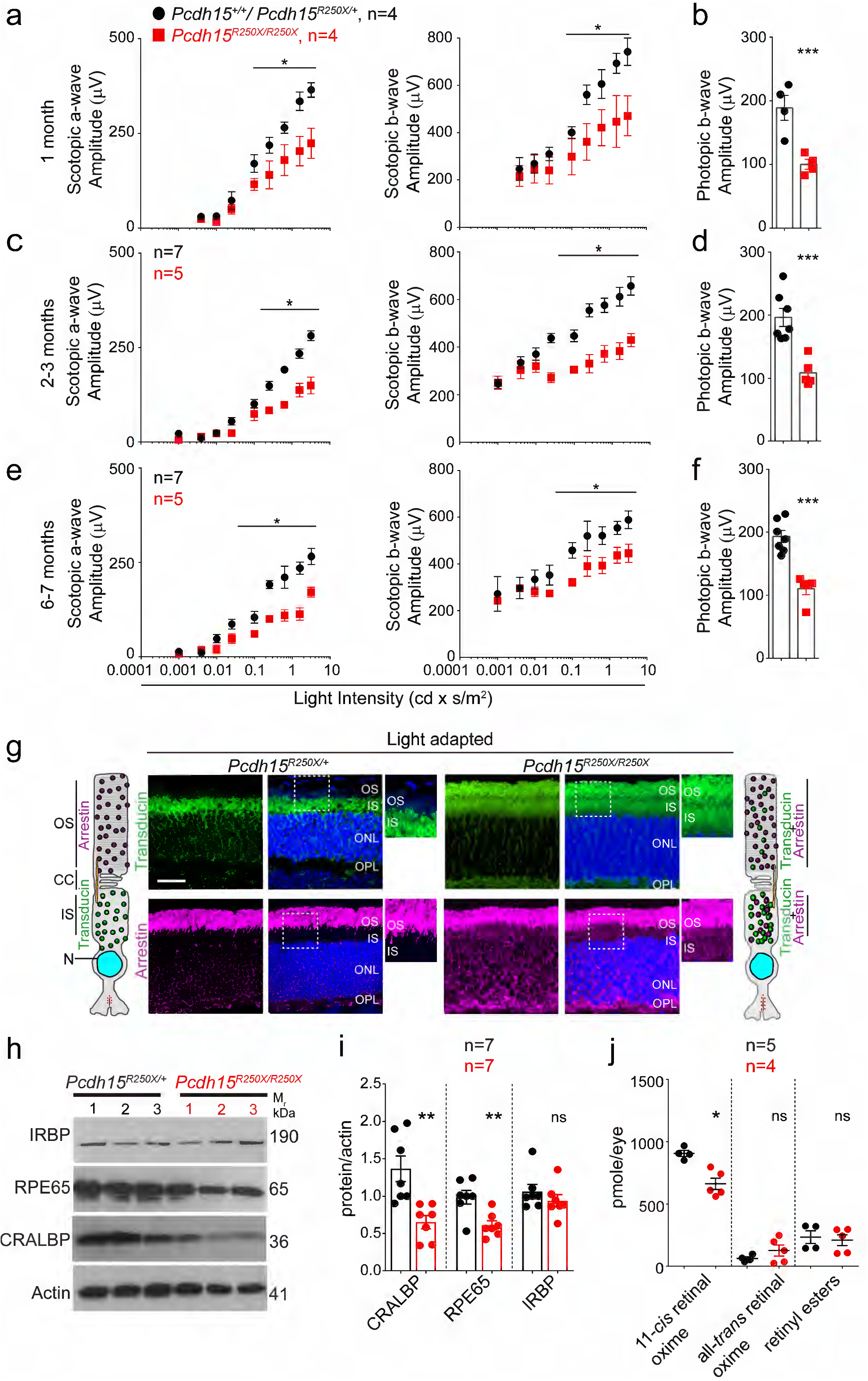
Loss of protocadherin-15 leads to visual dysfunction and aberrant localization of key proteins involved in the phototransdcution cascade and retinoid cycle. **a** Quantification of scotopic (dark adapted) responses from littermate control (*Pcdh15^+/+^* or *Pcdh15^R250X/+^*) and mutant (*Pcdh15^R250X/R250X^*) mice at one-month of age revealed progressive loss of both a- (*left* panels) and b-wave (*right* panels) amplitudes in mutant mice. Representative ERG waveforms are shown in **Extended data 3b**. **b** Quantification of photopic (light adapted) b-wave indicates decline of cone photoreceptor function in mutant mice. **c-f** Quantification of scotopic ERG amplitudes **c, e** and photopic ERG amplitudes **d, f** at indicated ages shows sustained decline in amplitudes over time in *Pcdh15* mutant mice. **g** Representative confocal micrographs of light adapted retinae shows mislocalization of phototransduction cascade proteins, arrestin and transducin, to both the inner segment (IS) and outer segment (OS) in mutant mice (*right* panels). In control mice, transducin is correctly localized to the IS and arrestin is to the OS (*left* panels). A schematic of the localization of arrestin and transducin in control and mutant mice is also shown. Scale bar – 20 μm. ONL – outer nuclear layer, OPL – outer plexiform layer. **h, i** Immunoblot of proteins involved in the visual retinoid cycle shows reduced quantities of RPE65 and CRALBP but not IRBP, quantified in **i**. **j** Quantification of indicated retinoid species from control and mutant mice shows reduced quantities of 11-*cis* retinal oxime. Data presented as mean ± SEM. Each data point represents an individual mouse. Data presented as mean ± SEM. Student unpaired *t*-test, *p*<0.05 (*).

### Mechanisms contributing to ERG defects due to protocadherin absence

We hypothesized that the functional deficits, reflected by abnormal ERG findings, without structural impairment of the retina might result from deficits in the phototransduction cascade or the visual retinoid cycle. The phototransduction cascade mediates the transduction of light into neuronal signals, while the visual retinoid cycle regenerates a key chromophore, 11-*cis* retinal. The 11-*cis* retinal complexes with opsin to form rhodopsin. Absorption of a single photon by 11-*cis* retinal leads to its photo-isomerization to all-*trans* retinal within femtoseconds (Nogly, Weinert et al., 2018), thus activating opsin and initiating the phototransduction cascade. Consequently, there is decoupling of opsin and all-*trans* retinal. All-*trans* retinal must be re-isomerized to 11-*cis* retinal to form rhodopsin again. These enzymatic steps occur in the retinal pigment epithelium (RPE)(Saari, 2000, Travis, Golczak et al., 2007, Wald & Brown, 1956). The rod outer segments (OS) are exquisitely adapted for light transduction. Phototransduction proteins are generated in the photoreceptor cell body and delivered to the outer segment (OS) via the inner segment (IS) and connecting cilium. Under photopic conditions (daylight), arrestin translocates from IS of the photoreceptors to the OS, to desensitize the opsin. Conversely, transducin translocates from OS to IS of the photoreceptors allowing arrestin to bind to opsin (Arshavsky, Lamb et al., 2002, Burns & Baylor, 2001). We found significant mislocalization of both arrestin and transducin to the photoreceptor IS and OS in *Pcdh15^R250X^* mutant mice under light-adapted conditions, whereas transducin was correctly localized only to the IS and arrestin only to the OS in control mouse retinae (**Fig. 2g, Fig. S4c, d**). In dark adapted conditions, arrestin was correctly localized to the IS and transducin to the OS in both mutant and control mice (**Fig. S4a, c-d**). Finally, opsin was correctly localized only in the OS under both dark- and light-adapted conditions (**Fig. S4b, e**), indicating that protocadherin-15 is essential for rapid shuttling of proteins from IS to OS and vice-versa in response to adaptation to light.

Next, we assessed the levels of crucial retinoid cycle proteins such as RPE65, an essential isomerase which catalyzes the conversion of all-*trans* retinyl ester to 11-*cis* retinol (Jin, Li et al., 2005, Moiseyev, Chen et al., 2005, Redmond, Poliakov et al., 2005), CRALBP (cellular retinaldehyde–binding protein), a key retinoid transporter, and IRBP (interphotoreceptor retinoid binding protein). These studies were rationalized based on the findings that protocadherin-15 is a binding partner of myosin VIIA(Senften et al., 2006), which also interacts with RPE65 (Lopes, Gibbs et al., 2011). Similar to protocadherin-15, pathogenic variants of myosin VIIA also cause USH1 (Jacobson, Cideciyan et al., 2011, Weston, Kelley et al., 1996). Intriguingly, immunoblotting revealed significantly reduced quantities of RPE65 and CRALBP, but not IRBP in *Pcdh15^R250X^* mutant mice (**Fig. 2h, i**). Next, we quantified the absolute retinoid levels within the eyes, and as compared to controls found reduced levels of retinoids, particularly 11-*cis* oxime, in *Pcdh15^R250X^* mutant mice (**Fig. 2j**). Together, our data indicate that loss of protocadherin-15 in the retina leads to aberrant translocation of proteins involved in the phototransduction cascade and reduced levels of key retinoids and enzymes involved in the visual retinoid cycle.

### Pre-clinical administration of exogenous retinoids

We hypothesized that low levels of retinoids in the mutant mice could be overcome by providing exogenous retinoids, thus rescuing the ERG deficits (Palczewski, 2010, Sethna, Scott et al., 2020). To test this hypothesis, we first performed baseline ERGs on 2-3-month-old control and *Pcdh15^R250X^* mutant mice. One week later, *Pcdh15^R250X^* mutant mice were injected intraperitoneally (IP) with 9-*cis* retinal, an analog of naturally occurring 11-*cis* retinal. Control mice were injected with vehicle. ERGs were performed the next day after overnight dark adaptation. Remarkably, a single treatment of *Pcdh15^R250X^* mutant mice with 9-*cis* retinal was sufficient to increase their ERG amplitudes to levels comparable to those in vehicle-injected controls (**Fig. 3a-b**). Similarly, we also observed an improvement in cone function to levels of the vehicle injected control mice (**Fig. 3c**). The b- to a-wave ratio was consistent with vehicle injected control mice or baseline *Pcdh15^R250X^* mutant mice (data not shown), suggesting a proportional increase in photoreceptor function after retinoid therapy.

**Figure 3:**
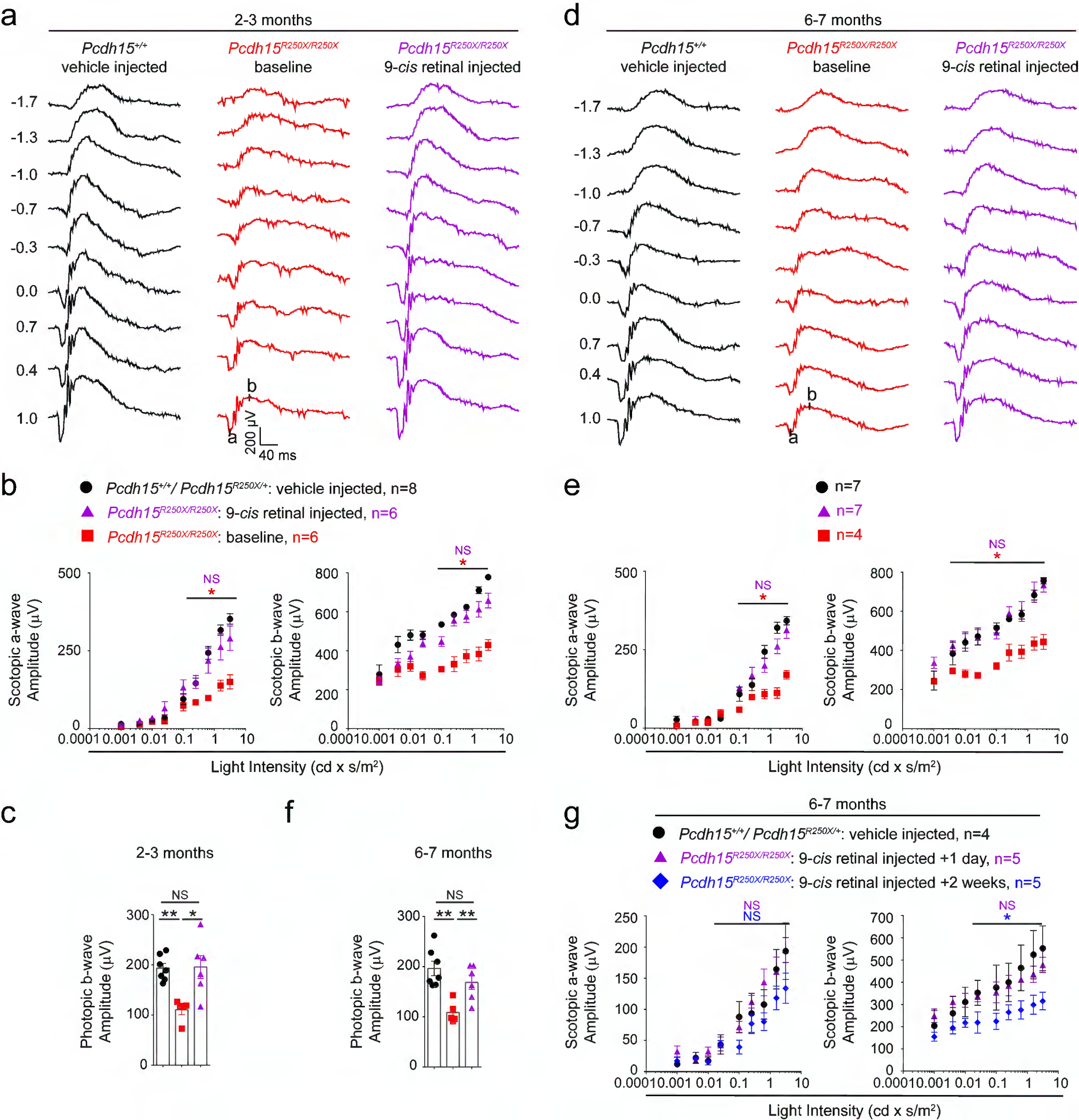
Exogenous 9-*cis* retinal rescues ERG deficits in young and old mutant mice. **a, d** Representative scotopic ERG traces from young (2-3 months) **a** and old (6-7 months) **d** 9-*cis* retinal injected *Pcdh15* mutant mice (*right* panels) show waveforms comparable to vehicle injected control mice (*left* panels). Same *Pcdh15^R250X^* mutant mice assessed 1 week prior to 9-*cis* retinal injection and ERG assessment show significantly reduced waveforms (*central* panels, baseline). **b, e** a- (*left* panel) and b-wave (*right* panel) quantification of scotopic ERG amplitudes shown in **a** and **d**, respectively. **c, f** Quantification of photopic b-wave for the denoted mice shows 9-*cis* retinal also improved cone-mediated function of mutant mice. **f** a- (*left* panel) and b-wave (*right* panel) quantification of scotopic ERG amplitudes for a different cohort of 6-7-months-old mice shows that 2 weeks after 9-*cis* retinal injection in mutant mice, the efficacy start to wane. Data presented as mean ± SEM. One-way ANOVA and Bonferroni *post hoc* test, *p*<0.05 (*) or <0.001 (***). NS – not significant.

Next, to assess the impact of exogenous retinoids in aged animals, we performed similar experiments using 6-7-month-old mice. We found a comparable increase in functional activity with a single IP injection of 9-*cis* retinal in mutant mice as compared to the same cohort of mutant mice assessed one week earlier (baseline *Pcdh15^R250X^* mutant mice). The ERG amplitudes of 9-*cis* retinal-injected *Pcdh15^R250X^* mutant mice were nearly indistinguishable from those of vehicle-injected control mice (**Fig. 3d-f**). Finally, in a separate cohort of 6-7-month-old mutant mice we assessed the longevity of the retinoid-mediated improvement. We found that by 2 weeks after 9-*cis* retinal treatment, the ERG improvement was waving off (**Fig. 3g**). Taken together, our data supports a role for visual retinoid cycle dysfunction in *Pcdh15^R250X^* mutant mice and provides a starting point to investigate the possibility of therapeutically boosting visual function in USH1F patients.

The spectrum and longitudinal ophthalmic phenotypes of USH1F patients homozygous for the p.Arg245* variant (or compound heterozygotes) consisted of a rod-cone dystrophy and are relatively uniform across this cohort of patients. They include an onset of symptoms such as night vision difficulties and visual field deficits in the first or early second decade, the presence of macular atrophy with reduction in central visual acuity by the third decade and, subsequently, the progressive constriction of visual fields resulting in tunnel vision between the third and fifth decades of life. Progressive posterior subcapsular cataract and optic nerve head atrophy are also frequent manifestations of this *USH1F* genotype.

*Pcdh15^R250X^* mutant mice have a retinal dysfunction as early as one month after birth. Our data indicate that protocadherin-15 has a dual role in photoreceptors and the RPE. First, at the junction of the photoreceptor IS and OS, where protocadherin-15 is localized (Reiners, Märker et al., 2005), the loss of protocadherin-15 leads to disrupted shuttling of arrestin and transducin under light-adapted conditions. Second, within the RPE, loss of protocadherin-15 is associated with lower levels of two key visual retinoid cycle enzymes, CRALBP and RPE65. Reduced levels of RPE65 were reported in *Myo7a* knockout mice (Lopes et al., 2011). Further, CRALBP facilitates the transport of 11-*cis* retinal between the RPE and the photoreceptor OS (Saari, Nawrot et al., 2001). Hence, reduced levels of RPE65 and CRALBP lead to delayed and reduced regeneration and transport of 11-*cis* retinal to the photoreceptor OS and hence we observed a concordant reduction in levels of 11-*cis* retinal oxime. Our data provide a plausible explanation for reduced ERG amplitudes without gross retinal degeneration in *Pcdh15^R250X^* mutants, suggesting this may also be the case for other *Pcdh15* mutant mice (Haywood-Watson et al., 2006a, Libby & Steel, 2001, Liu, Bulgakov et al., 2007, Peng, Zallocchi et al., 2011).

Unlike the typical human ocular manifestations of USH1, which have severe retinal degeneration, our mouse model has much less severe pathophysiology. This is consistent with reported ocular phenotypes of other USH mouse models (Haywood-Watson et al., 2006a, Jacobson et al., 2008, Liu et al., 2007, Liu, Udovichenko et al., 1999, Williams, Aleman et al., 2009). The discordance between retinal pathologies in humans and mice may be attributed to the structural differences in their photoreceptors, particularly the presence of calyceal process in humans, monkeys, and frog, but not in rodents (Sahly, Dufour et al., 2012, Schietroma, Parain et al., 2017), light exposure (Lopes et al., 2011), genetic backgrounds (Trouillet, Dubus et al., 2018), or environmental factors. Some of these possible factors are supported by recent observations of photoreceptor degeneration in *Ush1* frog models that have calyceal processes (Schietroma et al., 2017), as well as in mice on an albino background (Trouillet et al., 2018).

In conclusion, documenting the natural history and degree of clinical variability of ocular phenotype in human and animal models is pivotal for evaluating the efficacy of potential therapeutics in future clinical trials. Our longitudinal USH1F patient ocular data shows that significant vision and photoreceptors are preserved until the third decade of life, providing a long therapeutic window. Our results with an 11-*cis* retinal analog, 9-*cis* retinal, raises the possibility that longer lasting analogs such as 9-*cis* retinyl acetate, which has an excellent safety profile (Koenekoop, Sui et al., 2014, Scholl, Moore et al., 2015) or a synthetic version of 11-*cis* β-carotene, whose capsule formulation is already approved by the United States Food and Drug Administration (Rotenstreich, Belkin et al., 2013), could preserve vision in USH1F Usher syndrome patients. Furthermore, in mouse models lacking key visual cycle enzyme RPE65 or with the one of the most common variants of opsin causing RP (p.Pro23His), administration of retinoids has shown to preserve the photoreceptor morphology or proper folding of opsin to a greater extent (Maeda, Maeda et al., 2009, Noorwez, Malhotra et al., 2004), and thus might also extend the life of functional photoreceptors in USH1 patients. Based on our pre-clinical data in mouse and prior human trials, a clinical trial in USH1F patients may show benefit if the retinoid is administered early in life.

## METHODS

### Patient assessment

The records of 13 patients were reviewed under the National Eye Institute, National Institutes of Health protocol 08-EI-N014. Informed consents were obtained from the patients to conduct this research. Eleven were homozygous for the p.Arg245* founder variant was associated with the majority of Usher syndrome type 1 of the Ashkenazi Jews in our study and 2 subjects who were compound heterozygous with one p.Arg245* variant and a second pathogenic variant of *PCDH15* in trans. Data included demographic information, age of onset of visual symptoms, date of ophthalmic exams and reason for visit, best corrected visual acuity (BCVA), visual fields, presence and type of lenticular opacities, fundus exam findings, Optical Coherence Retinography when available and electroretinography. The horizontal diameter of the V4e, I4e, and I1e isopters on Goldmann Visual Fields were measured. Five patients were seen at the NIH Clinical Center under protocol 05-EI-0096 and three had follow-up visits over a thirty-year period. These patients underwent a complete ophthalmologic examination including best-corrected visual acuity (BCVA) with manifest refraction, biomicroscopy and photography of lens opacity, if present, and visual field evaluation by Goldmann kinetic perimetry (GVF). Dilated ophthalmoscopic examination was performed after instillation of phenylephrine 2.5% and tropicamide 1%. Digital photography of the retinal fundus was performed. Snellen visual acuity was measured using ETDRS charts. In patients whose visual acuity was reduced to a degree preventing them from reading the chart, the ability to recognize hand motion (HM) or perceive light (LP) was documented. The presence or absence of cystoid macular changes and/or atrophic pigmentary macular changes were assessed by ophthalmoscopy and/or macular photography.

### Generating *Pcdh15^R250X^* mice

*Pcdh15^R250X^* mice were generated by the Cincinnati Children’s Hospital Genetics Core using CRISPR/Cas9 technology and then transferred to the University of Maryland School of Medicine (UMSOM) facilities. In addition to the desired mutation (in red) that changes the codon of R250 from CGA to TGA, silent mutations (in green) were introduced to create a Hae II restriction endonuclease site (underlined) which is used to facilitate genotyping as well as to prevent recutting by Cas9 nuclease.

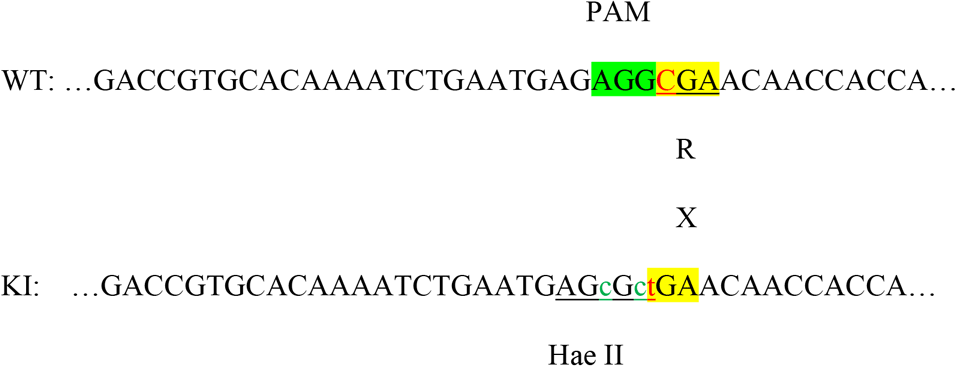

Heterozygous founder mice were bred with wild type (WT) C57BL/6J mice and the colony was further expanded on the C57BL/6J background. Mice are genotyped using primers VS4576: TTCACCTTCCATTCCCCCAAC and VS4577: CTTACCGGAGTCCTCAGTTCAGG, which generates a 343 bp amplicon that was also Sanger sequenced. Mice were housed in a facility with 12 hours of light and 12 hours with the lights off. Mice were fed after weaning on a standard mouse diet and with water available *ad libitum*. We followed the ARRIVE guidelines for reporting animal research and studies were conducted in accordance with the ARVO Statement for the Use of Animals in Ophthalmic and Vision Research as well as the National Institutes of Health Guide for the Care and Use of Laboratory Animals. All animal procedures were approved by the UMSOM IACUC (Institutional Animal Care and Use Committees).

### Electroretinography (ERG) and Optical Coherence Tomography (OCT)

ERGs were recorded as previously described (Sethna, Chamakkala et al., 2016). Overnight dark-adapted mice were anesthetized with a combination of ketamine-xylazine (100 mg/kg and 10 mg/kg, respectively), followed by dilation of pupils with 1% Tropicamide. A gold loop wire electrode was placed on the cornea, a reference electrode was placed on the scalp under the skin and a ground electrode was placed under the skin near the tail. ERG waveforms were acquired using sequentially brighter stimuli (0.003962233 to 3.147314 cd × s/m^2^) with 5-60 sec intervals using the Diagnosys ColorDome Ganzfeld system (Diagnosys systems, Lowell, MA). Three to five waveforms per intensity were averaged. Photopic, cone-only, responses were acquired at a single bright flash (3.15 cd × s/m^2^) under a steady rod-suppressing field of 30 cd × s/m^2^, with 10 waves averaged. Waves were analyzed using inbuilt Espion software. For exogenous 9-*cis* retinal treatment, animals received intraperitoneal 0.25 mg 9-*cis* retinal (dissolved in 100% ethanol) in 200 μl of vehicle (sterile filtered 10% BSA in 0.9% NaCl solution) or vehicle only, in the dark. ERGs were performed as above. OCT was performed using Spectralis OCT (Heidelberg Engineering, Heidelberg, Germany). Mice were anesthetized and dilated as above. A custom designed plano-concave contact lens micro-M 2.00/5.00 (Cantor & Nissel Ltd, Northamptonshire, UK) was used to obtain cross sections of the entire retina.

### Immunohistochemistry and FM1-43 uptake

Mice (1 to 3-month-old) were dark adapted overnight and euthanized before light onset and eyes were enucleated following CO_2_ asphyxiation followed by cervical dislocation or exposed to normal room light for 2 hours after light onset and euthanized as above and processed as below. Dark adapted procedures were performed under very dim red light. Eyes were immediately fixed in Prefer fixative (Anatech LTD, Battle Creek, MI), paraffin embedded and stained using standard protocols (Sethna et al., 2016, Sethna & Finnemann, 2013). Briefly, 7 μm sections were deparaffinized, rehydrated in PBS, blocked and permeabilized with 10% normal goat serum/0.3% triton-X 100 for 2 hours at RT, and incubated overnight at 4°C with the indicated primary antibodies to transducin (1:100 dilution, #Sc-517057, Santa Cruz Biotechnology) and arrestin (clone C10C10, 1:25 dilution, kind gift from Drs. Paul Hargrave and Clay Smith, University of Florida). The following day, sections were incubated with Alexa fluor labeled goat secondary antibodies (1:250) and DAPI (Thermo Fisher Scientific, Waltham, MA) to label nuclei. Sections were scanned using the UMSOM core facility Nikon W1 spinning disk microscope and images were processed using FIJI software (Schindelin, Arganda-Carreras et al., 2012). To stain for protocadherin-15, dissected eyes were fixed in 4% PFA (Electron Microscopy Sciences, Hatfield, PA) and processed as above using a previously described custom antibody targeting the C-terminus of protocadherin-15 (PB303; 1:200) (Ahmed, Riazuddin et al., 2003a).

P60 temporal bones were fixed and processed for immunocytochemistry as previously described(Riazuddin, Belyantseva et al., 2012). The cochlear and vestibular sensory epithelia were isolated, fine dissected and permeabilized in 0.25% Triton X-100 for 1 hour and blocked with 10% normal goat serum in PBS for 1 hour. Tissue samples were probed overnight with antibodies against myosin VIIa or custom antibody targeting the N-terminus of antibody protocadherin-15 (HL5164; 1:200 dilution)(Ahmed et al., 2006b), and after three washes, were incubated with the secondary antibody for 45 mins at room temperature. Rhodamine phalloidin was used at a 1:250 dilution for F-actin labeling. All images were acquired using a LSM 700 laser scanning confocal microscope (Zeiss, Germany) using a 63×1.4 NA or 100×1.4 NA oil immersion objectives. Stacks of confocal images were acquired with a Z step of 0.5μm and processed using ImageJ software (National Institutes of Health).

Cochlear and vestibular explants were dissected at postnatal day 0 (P0) and cultured in a glass-bottom petri dish (MatTek, Ashland, MA). They were maintained in Dulbecco’s modified Eagle medium (DMEM) supplemented with 10% FBS (Thermo Fisher Scientific, Waltham, MA) for 2 days at 37 °C and 5% CO2. Explants were incubated for 10 sec with 3 μM FM1-43, washed three times with Hank’s balanced salt solution, and imaged live using a Zeiss LSM 700 scanning confocal microscope.

### Auditory Brainstem Response (ABR) measurements

Hearing thresholds of heterozygous and homozygous *Pcdh15^R250X^* mice at P16 (n = 5 each genotype) were evaluated by recording ABRs. All ABR recordings, including broadband clicks and tone-burst stimuli at three frequencies (8, 16, and 32 kHz), were performed using an auditory-evoked potential RZ6-based auditory workstation (Tucker-Davis Technologies) with a high frequency transducer. Maximum sound intensity tested was 100 dB SPL. TDT system III hardware and BioSigRZ software (Tucker Davis Technology) were used for stimulus presentation and response averaging.

### Vestibular testing

Exploratory tests were performed as previously described (Michel, Booth et al., 2017). Briefly, mice were placed individually in a new cage. A camera was placed on top of the cage to record movements of mice for 2 mins and tracked using ImageJ software. Tail hanging tests were performed as follow: Mice were held five centimeters above a tabletop. The test scores were given as following: normal behavior was demonstrated by a “reaching position”, with a score of 4, by the extension of limb and head forward and downward aiming to the tabletop. Mice with abnormal behavior, ranked with a score of 1, tried to climb towards the examiner’s hand, curling the body upward reaching with the head to the tail one time. Mice ranked with a score of 0, tried to climb towards the examiner’s hand, curling the body upward reaching with the head to the tail multiple times.

### Retinoid extraction and analysis

All procedures for retinoid extraction were performed under red safelights. Mice were euthanized with CO_2_, eyes enucleated, lens and vitreous removed, followed by freezing the eyecup on dry ice, which were stored at −80°C until retinoid extraction was performed. Mouse eyecups were homogenized in fresh hydroxylamine buffer (1 ml of 50 mM MOPS, 10 mM NH2OH, pH 6.5). 1 ml ethanol was added, samples were mixed and incubated (30 min in the dark at RT). Retinoids were extracted into hexane (2 × 4 ml), followed by solvent evaporation using a gentle stream of argon at 37°C. After reconstituting in 50 μl mobile phase, samples were separated on a 5 μm LiChrospher Si-60 (ES Industries, West Berlin, NJ, USA) normal-phase column on a Waters H-Class Acquity UPLC (Waters Corp., Milford, MA, USA) along with standards at a flow rate of 1.5 ml/min, following published methods(Landers & Olson, 1988). Retinaloxime standards were prepared from 11-*cis* retinal using published methods(Garwin & Saari, 2000) and retinyl palmitate for use as standard was from Sigma Aldrich Inc. (Saint Louis, MO, USA). Absorbance was monitored at 350 nm for retinaloximes and at 325 nm for retinyl esters. Peak areas were integrated and quantified using external calibration curves. Data were analyzed using Empower 3 software (Waters, Milford, MA, USA).

### Data analysis

Four to eight animals per time point/genotype/treatment for ERG analysis were used. One-way ANOVA with Tukey’s post-hoc test or Student’s *t*-test was used to compare control sample to test samples with the data presented as mean ± SEM. Differences with p < 0.05 were considered significant. Data were analyzed using GraphPad Prism (GraphPad Software, Inc., La Jolla, CA, USA).

### Contact for reagent and resource sharing

Further information and requests for resources and reagents should be directed to and will be fulfilled by the Lead Contact, Zubair M. Ahmed, PhD.

## Contributions

W.M.Z., A.J.G., and T.B.F. conceived and designed the human research study, while S.S. and Z.M.A. designed and conceived the *Pcdh15* mouse model project. S.S., W.M.Z., S.R., A.P.J.G., J.M.S., T.D., R.B.H., and C.C.B. carried out the experimental or clinical work and analyzed data. A.J.G., T.M.R., Sa.R., T.B.F., and Z.M.A. provided resources, supervised the experiments, and analyzed data. S.S., W.M.Z., T.B.F., and Z.M.A. wrote the manuscript. All authors read, edited, and approved the manuscript.

## Acknowledgments

We thank all the probands for participating in the natural history studies. We thank Dr. Ekaterina Tsilou for clinical assessments, Ms. Amy Turriff and Ms. Meira Meltzer for genetic counselling, Ms. Dimitria Gomes and Mr. Samuel Garmoe for technical assistance with mice and the UMSOM core facility for access to a Zeiss-710 confocal and Nikon W1 microscopes. We appreciate review of the manuscript by Drs. Wade Chien and Isabelle Roux. The natural history project at the National Eye Institute (NEI) and National Institute on Deafness and Other Communication Disorders (NIDCD) was supported (in part) by intramural funds to W.M.Z. and T.B.F (DC000039), respectively. Work at University of Maryland Baltimore was supported by Research Funds from Usher1F Collaborative Foundation award (Z.M.A.).

---

**Fig. S1:**
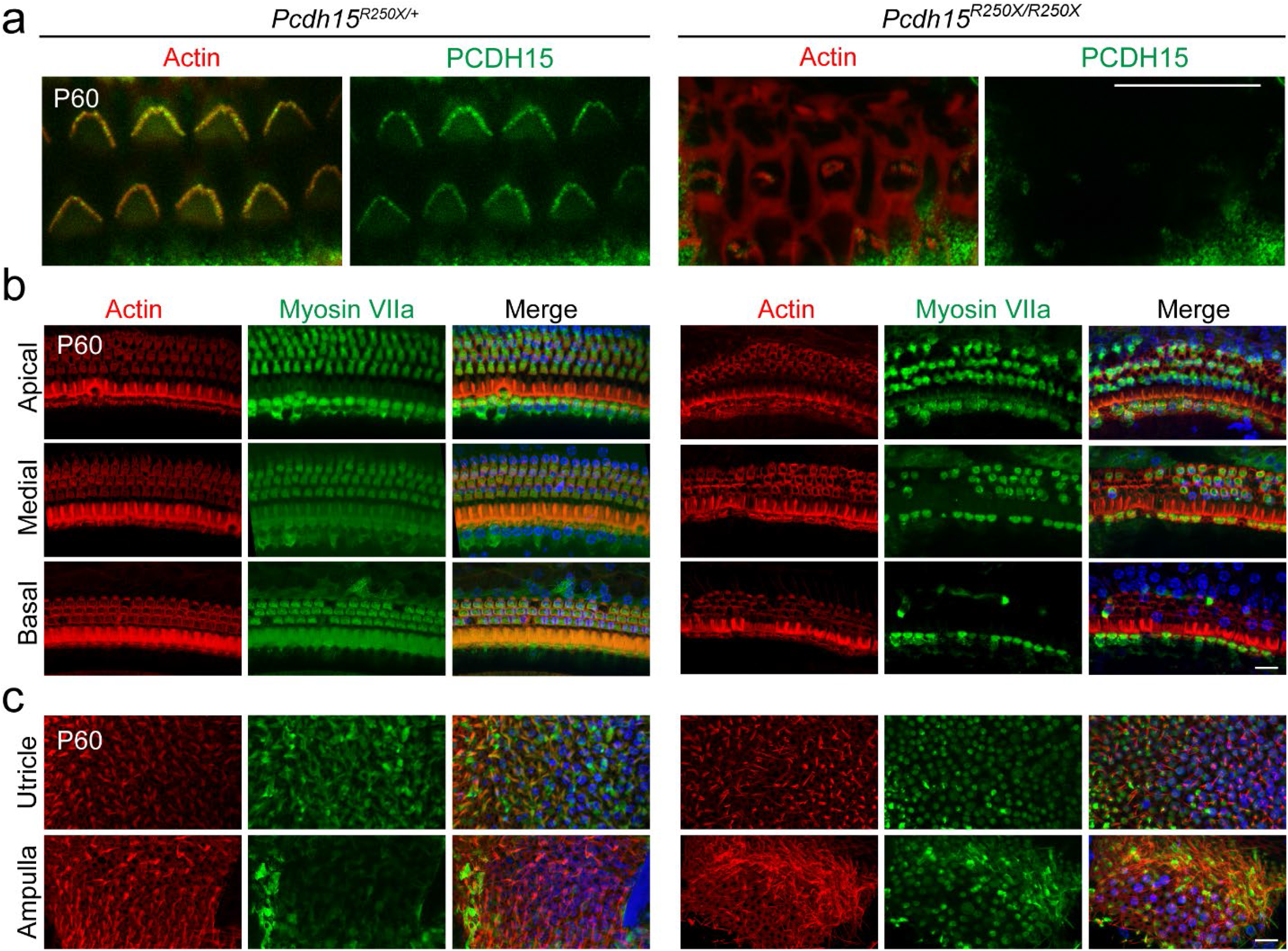
*Pcdh15^R250X^* mutant mice have degeneration of the organ of Corti. **a** Confocal micrographs of P60 organ of Corti of indicated genotype immunostained with protocadherin-15 (PCDH15) antibody HL5614 (green) and counterstained with phalloidin (red) confirms the loss of protocadherin-15 in *Pcdh15^R250X/^*^R250X^ mice. **b, c** Confocal micrographs of P60 organ of Corti **b** and vestibular end organs **c** of control (*Pcdh15^R250X/+^*) and mutant (*Pcdh15^R250X/R250X^*) mice immunostained with myosin VIIa antibody (green) and counterstained with phalloidin (red) and DAPI (blue), revealed degeneration of sensory hair cells. Medial and basal turns of the cochlea and utricle are shown. Scale bar, 10 μm.

**Fig. S2:**
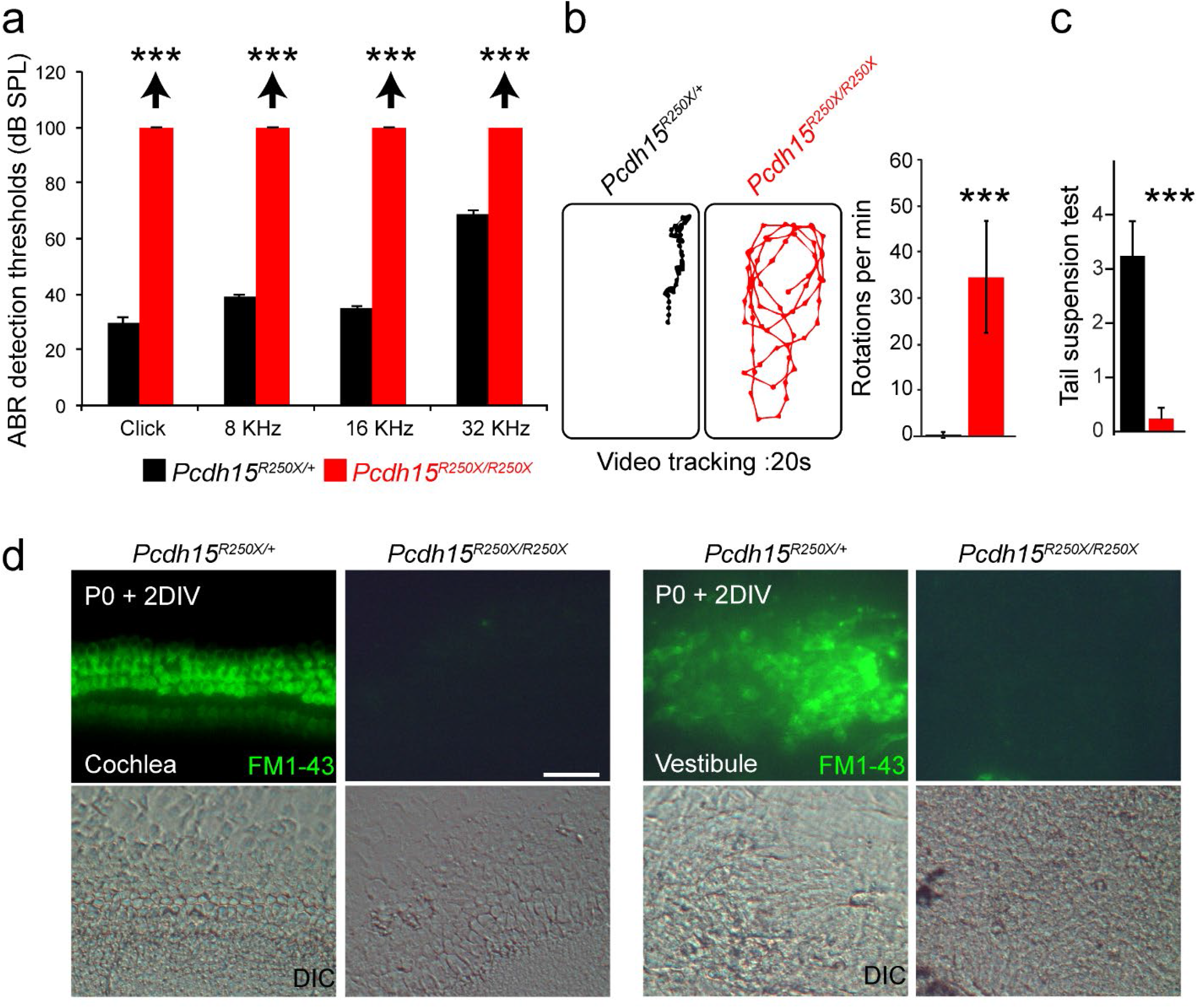
*Pcdh15^R250X^* mutant mice have profound hearing loss and severe vestibular system dysfunction. **a** Average thresholds of acoustic brainstem response (ABR) to broadband clicks and tone-pips with frequencies of 8 kHz, 16 kHz, and 32 kHz in control (*Pcdh15^R250X/+^*; black) and mutant (*Pcdh15^R250X/R250X^*; red) mice at P16-P30 (n = 4/ genotype), revealed no detectable hearing in *Pcdh15^R250X^* mutants. **b** Representative open‐field exploratory behavior of a 2-month old mouse for denoted genotypes shows increased circling behavior in mutant mice (*left* panels), quantified in *right* panel. (*n* = 3/*genotype*). **c** Average score of tail-suspension test in control and mutants at P16-30, further confirm vestibular areflexia in *Pcdh15^R250X^* mutant mice. All data are shown as Mean ± SEM. (***, *p*<0.001) using Student unpaired *t*-test. **d** Epifluorescence micrographs and corresponding DIC micrographs of control and *Pcdh15^R250X/R25X^* cultured organ of Corti and vestibular explants imaged after exposure to 3μM of FM1-43, a channel permeable dye, for 10 sec, revealed impaired mechanotransduction function in *Pcdh15^R250X^* mutant mice. The samples were dissected at P0 and kept two days *in vitro* (P0+ 2DIV). Scale bar: 20 μm.

**Fig. S3:**
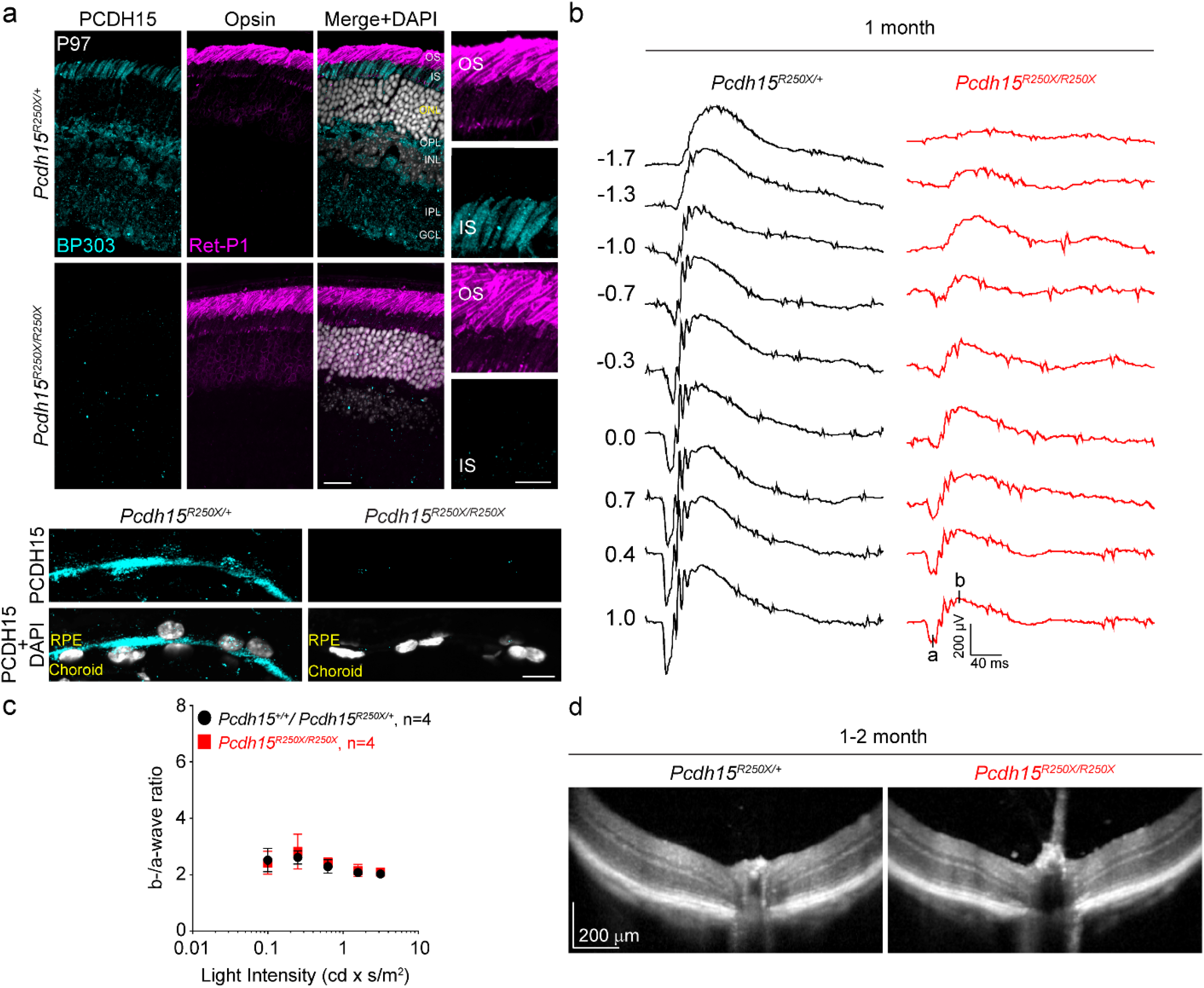
Validation of loss of protocadherin-15 in the retinae of *Pcdh15^R250X^* mutant mice, and associated eye phenotype. **a** Confocal micrographs of P97 mouse retinae from control and *Pcdh15^R250X^* mutants immunostained for protocadherin-15 (PCDH15, cyan, *left* panel) and opsin to label outer segments (OS, magenta, *center* panel) shows robust protocadherin 15 localized to the inner segments (IS), outer plexiform layer (OPL) and ganglion cell layer (GCL). Notably, protocadherin-15 immunoreactivity is absent in *Pcdh15^R250X^* mutant mice. *Inset* shows zoomed in image of the OS/ IS interface. DAPI to visualize nuclei in grey. Scale bar, 10 μm. **b** Representative ERG waveforms from 1-month old mice of the denoted genotypes shows normal wave architecture with significantly reduced waveforms for mutant mice. **c** b/a ratio for mice depicted in **b** shows the mutant mice have a ratio comparable to control mice. **d** OCT images from 1-month old mice of denoted genotypes shows no gross retinal degeneration in mutant mice. INL – inner nuclear layer; IPL – inner plexiform layer.

**Fig. S4:**
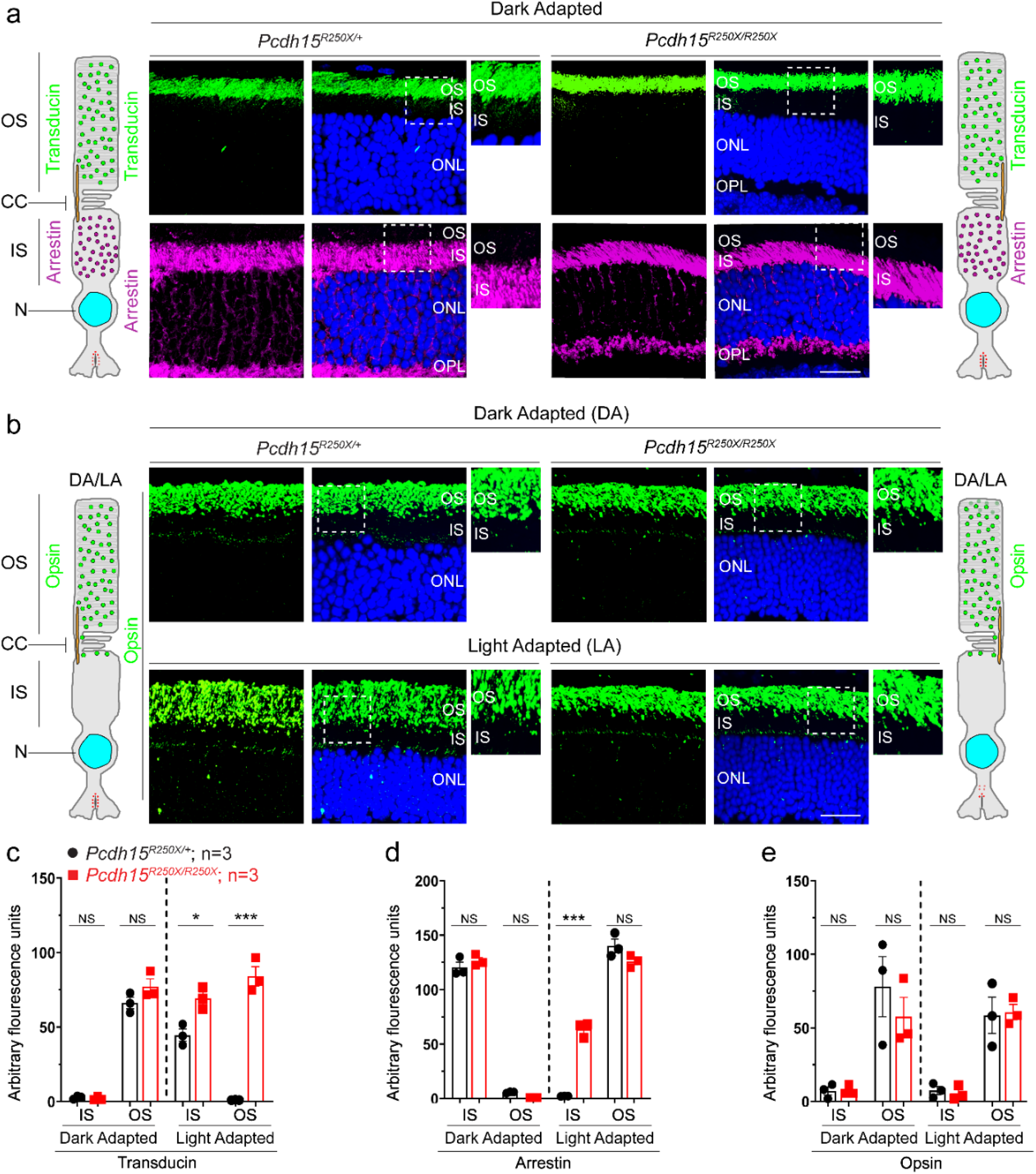
Loss of protocadherin-15 does not affect dark adapted localization of key phototransduction cascade proteins. **a** Representative confocal micrographs of dark adapted retinae show the correct localization of phototransduction cascade proteins, transducin to the outer segment (OS) and arrestin to the inner segment (IS), in both control and mutant mice. Schematic of the localization of arrestin and transducin in control and mutant mice is also shown. **b** Dark adapted (*top* panels) and light adapted (*bottom* panels) localization of opsin is not affected by loss of protocadherin-15. Schematic of the localization of opsin in control and mutant mice is also shown. Scale bar – 20 μm. ONL – outer nuclear layer, OPL – outer plexiform layer. **c-e** Quantification of denoted proteins in the inner segment (IS) or outer segment (OS) for panels in **a, b** and **Fig. 2g** shows mislocalization of transducin to the OS and arrestin to the IS in mutant mice. Student unpaired *t-test*, *p* <0.05 (*), *p* <0.001 (***), NS – not significant.

## REFERENCES

Ahmed ZM, Goodyear R, Riazuddin S, Lagziel A, Legan PK, Behra M, Burgess SM, Lilley KS, Wilcox ER, Griffith AJ, Frolenkov GI, Belyantseva IA, Richardson GP, Friedman TB (2006a) The tip-link antigen, a protein associated with the transduction complex of sensory hair cells, is protocadherin-15. J Neurosci 26: 7022–34

Ahmed ZM, Goodyear R, Riazuddin S, Lagziel A, Legan PK, Behra M, Burgess SM, Lilley KS, Wilcox ER, Riazuddin S, Griffith AJ, Frolenkov GI, Belyantseva IA, Richardson GP, Friedman TB (2006b) The Tip-Link Antigen, a Protein Associated with the Transduction Complex of Sensory Hair Cells, Is Protocadherin-15. The Journal of Neuroscience 26: 7022–7034

Ahmed ZM, Riazuddin S, Ahmad J, Bernstein SL, Guo Y, Sabar MF, Sieving P, Riazuddin S, Griffith AJ, Friedman TB, Belyantseva IA, Wilcox ER (2003a) PCDH15 is expressed in the neurosensory epithelium of the eye and ear and mutant alleles are responsible for both USH1F and DFNB23. Human Molecular Genetics 12: 3215–3223

Ahmed ZM, Riazuddin S, Bernstein SL, Ahmed Z, Khan S, Griffith AJ, Morell RJ, Friedman TB, Riazuddin S, Wilcox ER (2001a) Mutations of the protocadherin gene PCDH15 cause Usher syndrome type 1F. American journal of human genetics 69: 25–34

Ahmed ZM, Riazuddin S, Bernstein SL, Ahmed Z, Khan S, Griffith AJ, Morell RJ, Friedman TB, Wilcox ER (2001b) Mutations of the protocadherin gene PCDH15 cause Usher syndrome type 1F. Am J Hum Genet 69: 25–34

Ahmed ZM, Riazuddin S, Riazuddin S, Wilcox ER (2003b) The molecular genetics of Usher syndrome. Clin Genet 63: 431–44

Alagramam KN, Goodyear RJ, Geng R, Furness DN, van Aken AFJ, Marcotti W, Kros CJ, Richardson GP (2011) Mutations in protocadherin 15 and cadherin 23 affect tip links and mechanotransduction in mammalian sensory hair cells. PloS one 6: e19183–e19183

Arshavsky VY, Lamb TD, Pugh EN (2002) G Proteins and Phototransduction. Annual Review of Physiology 64: 153–187

Ben-Yosef T, Ness SL, Madeo AC, Bar-Lev A, Wolfman JH, Ahmed ZM, Desnick RJ, Willner JP, Avraham KB, Ostrer H, Oddoux C, Griffith AJ, Friedman TB (2003) A Mutation of PCDH15 among Ashkenazi Jews with the Type 1 Usher Syndrome. New England Journal of Medicine 348: 1664–1670

Brownstein Z, Ben-Yosef T, Dagan O, Frydman M, Abeliovich D, Sagi M, Abraham FA, Taitelbaum-Swead R, Shohat M, Hildesheimer M, Friedman TB, Avraham KB (2004a) The R245X mutation of PCDH15 in Ashkenazi Jewish children diagnosed with nonsyndromic hearing loss foreshadows retinitis pigmentosa. Pediatr Res 55: 995–1000

Brownstein Z, Ben-Yosef T, Dagan O, Frydman M, Abeliovich D, Sagi M, Abraham FA, Taitelbaum-Swead R, Shohat M, Hildesheimer M, Friedman TB, Avraham KB (2004b) The R245X Mutation of PCDH15 in Ashkenazi Jewish Children Diagnosed with Nonsyndromic Hearing Loss Foreshadows Retinitis Pigmentosa. Pediatric Research 55: 995

Burns ME, Baylor DA (2001) Activation, Deactivation, and Adaptation in Vertebrate Photoreceptor Cells. Annual Review of Neuroscience 24: 779–805

Cong L, Ran FA, Cox D, Lin S, Barretto R, Habib N, Hsu PD, Wu X, Jiang W, Marraffini LA, Zhang F (2013) Multiplex Genome Engineering Using CRISPR/Cas Systems. Science (New York, NY) 339: 819–823

Garwin GG, Saari JC (2000) [19] High-performance liquid chromatography analysis of visual cycle retinoids. In Methods in Enzymology, pp 313–324. Academic Press

Haywood-Watson IIRJL, Ahmed ZM, Kjellstrom S, Bush RA, Takada Y, Hampton LL, Battey JF, Sieving PA, Friedman TB (2006a) Ames Waltzer Deaf Mice Have Reduced Electroretinogram Amplitudes and Complex Alternative Splicing of Pcdh15 Transcripts. Investigative Ophthalmology & Visual Science 47: 3074–3084

Haywood-Watson RJ, 2nd, Ahmed ZM, Kjellstrom S, Bush RA, Takada Y, Hampton LL, Battey JF, Sieving PA, Friedman TB (2006b) Ames Waltzer deaf mice have reduced electroretinogram amplitudes and complex alternative splicing of Pcdh15 transcripts. Invest Ophthalmol Vis Sci 47: 3074–84

Jacobson SG, Cideciyan AV, Aleman TS, Sumaroka A, Roman AJ, Gardner LM, Prosser HM, Mishra M, Bech-Hansen NT, Herrera W, Schwartz SB, Liu X-Z, Kimberling WJ, Steel KP, Williams DS (2008) Usher syndromes due to MYO7A, PCDH15, USH2A or GPR98 mutations share retinal disease mechanism. Human Molecular Genetics 17: 2405–2415

Jacobson SG, Cideciyan AV, Gibbs D, Sumaroka A, Roman AJ, Aleman TS, Schwartz SB, Olivares MB, Russell RC, Steinberg JD, Kenna MA, Kimberling WJ, Rehm HL, Williams DS (2011) Retinal Disease Course in Usher Syndrome 1B Due to MYO7A Mutations. Investigative Ophthalmology & Visual Science 52: 7924–7936

Jin M, Li S, Moghrabi WN, Sun H, Travis GH (2005) Rpe65 Is the Retinoid Isomerase in Bovine Retinal Pigment Epithelium. Cell 122: 449–459

Kazmierczak P, Sakaguchi H, Tokita J, Wilson-Kubalek EM, Milligan RA, Muller U, Kachar B (2007) Cadherin 23 and protocadherin 15 interact to form tip-link filaments in sensory hair cells. Nature 449: 87–91

Koenekoop RK, Sui R, Sallum J, van den Born LI, Ajlan R, Khan A, den Hollander AI, Cremers FPM, Mendola JD, Bittner AK, Dagnelie G, Schuchard RA, Saperstein DA (2014) Oral 9-cis retinoid for childhood blindness due to Leber congenital amaurosis caused by RPE65 or LRAT mutations: an open-label phase 1b trial. The Lancet 384: 1513–1520

Landers GM, Olson JA (1988) Rapid, simultaneous determination of isomers of retinal, retinal oxime and retinol by high-performance liquid chromatography. Journal of Chromatography A 438: 383–392

Libby RT, Steel KP (2001) Electroretinographic Anomalies in Mice with Mutations in Myo7a, the Gene Involved in Human Usher Syndrome Type 1B. Investigative Ophthalmology & Visual Science 42: 770–778

Liu X, Bulgakov OV, Darrow KN, Pawlyk B, Adamian M, Liberman MC, Li T (2007) Usherin is required for maintenance of retinal photoreceptors and normal development of cochlear hair cells. Proceedings of the National Academy of Sciences 104: 4413–4418

Liu X, Udovichenko IP, Brown SDM, Steel KP, Williams DS (1999) Myosin VIIa Participates in Opsin Transport through The Photoreceptor Cilium. The Journal of Neuroscience 19: 6267–6274

Lopes VS, Gibbs D, Libby RT, Aleman TS, Welch DL, Lillo C, Jacobson SG, Radu RA, Steel KP, Williams DS (2011) The Usher 1B protein, MYO7A, is required for normal localization and function of the visual retinoid cycle enzyme, RPE65. Human Molecular Genetics 20: 2560–2570

Maeda T, Maeda A, Casadesus G, Palczewski K, Margaron P (2009) Evaluation of 9-cis-Retinyl Acetate Therapy in Rpe65 −/− Mice. Investigative Ophthalmology & Visual Science 50: 4368–4378

Michel V, Booth KT, Patni P, Cortese M, Azaiez H, Bahloul A, Kahrizi K, Labbé M, Emptoz A, Lelli A, Dégardin J, Dupont T, Aghaie A, Oficjalska-Pham D, Picaud S, Najmabadi H, Smith RJ, Bowl MR, Brown SD, Avan P et al. (2017) CIB2, defective in isolated deafness, is key for auditory hair cell mechanotransduction and survival. EMBO Mol Med 9: 1711–1731

Moiseyev G, Chen Y, Takahashi Y, Wu BX, Ma J-x (2005) RPE65 is the isomerohydrolase in the retinoid visual cycle. Proceedings of the National Academy of Sciences of the United States of America 102: 12413–12418

Nogly P, Weinert T, James D, Carbajo S, Ozerov D, Furrer A, Gashi D, Borin V, Skopintsev P, Jaeger K, Nass K, Båth P, Bosman R, Koglin J, Seaberg M, Lane T, Kekilli D, Brünle S, Tanaka T, Wu W et al. (2018) Retinal isomerization in bacteriorhodopsin captured by a femtosecond x-ray laser. Science 361: eaat0094

Noorwez SM, Malhotra R, McDowell JH, Smith KA, Krebs MP, Kaushal S (2004) Retinoids assist the cellular folding of the autosomal dominant retinitis pigmentosa opsin mutant P23H. J Biol Chem 279: 16278–84

Palczewski K (2010) Retinoids for Treatment of Retinal Diseases. Trends in pharmacological sciences 31: 284–295

Peng Y-W, Zallocchi M, Wang W-M, Delimont D, Cosgrove D (2011) Moderate Light-Induced Degeneration of Rod Photoreceptors with Delayed Transducin Translocation in shaker1 Mice. Investigative Ophthalmology & Visual Science 52: 6421–6427

Pennings RJ, Damen GW, Snik AF, Hoefsloot L, Cremers CW, Mylanus EA (2006) Audiologic performance and benefit of cochlear implantation in Usher syndrome type I. Laryngoscope 116: 717–22

Redmond TM, Poliakov E, Yu S, Tsai J-Y, Lu Z, Gentleman S (2005) Mutation of key residues of RPE65 abolishes its enzymatic role as isomerohydrolase in the visual cycle. Proceedings of the National Academy of Sciences of the United States of America 102: 13658

Reiners J, Märker T, Jürgens K, Reidel B, Wolfrum U (2005) Photoreceptor expression of the Usher syndrome type 1 protein protocadherin 15 (USH1F) and its interaction with the scaffold protein harmonin (USH1C). Mol Vis 11: 347–55

Reiners J, van Wijk E, Marker T, Zimmermann U, Jurgens K, te Brinke H, Overlack N, Roepman R, Knipper M, Kremer H, Wolfrum U (2005) Scaffold protein harmonin (USH1C) provides molecular links between Usher syndrome type 1 and type 2. Hum Mol Genet 14: 3933–43

Riazuddin S, Belyantseva IA, Giese A, Lee K, Indzhykulian AA, Nandamuri SP, Yousaf R, Sinha GP, Lee S, Terrell D, Hegde RS, Ali RA, Anwar S, Andrade-Elizondo PB, Sirmaci A, Parise LV, Basit S, Wali A, Ayub M, Ansar M et al. (2012) Mutations in CIB2, a calcium and integrin binding protein, cause Usher syndrome type 1J and nonsyndromic deafness DFNB48. Nature genetics 44: 1265–1271

Rotenstreich Y, Belkin M, Sadetzki S, et al. (2013) Treatment with 9-cis β-carotene–rich powder in patients with retinitis pigmentosa : A randomized crossover trial. JAMA Ophthalmology 131: 985–992

Saari JC (2000) Biochemistry of Visual Pigment Regeneration The Friedenwald Lecture. Investigative Ophthalmology & Visual Science 41: 337–348

Saari JC, Nawrot M, Kennedy BN, Garwin GG, Hurley JB, Huang J, Possin DE, Crabb JW (2001) Visual Cycle Impairment in Cellular Retinaldehyde Binding Protein (CRALBP) Knockout Mice Results in Delayed Dark Adaptation. Neuron 29: 739–748

Sahly I, Dufour E, Schietroma C, Michel V, Bahloul A, Perfettini I, Pepermans E, Estivalet A, Carette D, Aghaie A, Ebermann I, Lelli A, Iribarne M, Hardelin J-P, Weil D, Sahel J-A, El-Amraoui A, Petit C (2012) Localization of Usher 1 proteins to the photoreceptor calyceal processes, which are absent from mice. The Journal of Cell Biology 199: 381–399

Schietroma C, Parain K, Estivalet A, Aghaie A, Boutet de Monvel J, Picaud S, Sahel J-A, Perron M, El-Amraoui A, Petit C (2017) Usher syndrome type 1–associated cadherins shape the photoreceptor outer segment. The Journal of Cell Biology

Schindelin J, Arganda-Carreras I, Frise E, Kaynig V, Longair M, Pietzsch T, Preibisch S, Rueden C, Saalfeld S, Schmid B, Tinevez J-Y, White DJ, Hartenstein V, Eliceiri K, Tomancak P, Cardona A (2012) Fiji: an open-source platform for biological-image analysis. Nature Methods 9: 676

Scholl HPN, Moore AT, Koenekoop RK, Wen Y, Fishman GA, van den Born LI, Bittner A, Bowles K, Fletcher EC, Collison FT, Dagnelie G, Degli Eposti S, Michaelides M, Saperstein DA, Schuchard RA, Barnes C, Zein W, Zobor D, Birch DG, Mendola JD et al. (2015) Safety and Proof-of-Concept Study of Oral QLT091001 in Retinitis Pigmentosa Due to Inherited Deficiencies of Retinal Pigment Epithelial 65 Protein (RPE65) or Lecithin:Retinol Acyltransferase (LRAT). PLOS ONE 10: e0143846

Senften M, Schwander M, Kazmierczak P, Lillo C, Shin J-B, Hasson T, Géléoc GSG, Gillespie PG, Williams D, Holt JR, Müller U (2006) Physical and Functional Interaction between Protocadherin 15 and Myosin VIIa in Mechanosensory Hair Cells. The Journal of Neuroscience 26: 2060

Sethna S, Chamakkala T, Gu X, Thompson TC, Cao G, Elliott MH, Finnemann SC (2016) Regulation of Phagolysosomal Digestion by Caveolin-1 of the Retinal Pigment Epithelium Is Essential for Vision. Journal of Biological Chemistry 291: 6494–6506

Sethna S, Finnemann SC (2013) Analysis of Photoreceptor Rod Outer Segment Phagocytosis by RPE Cells In Situ. In Retinal Degeneration: Methods and Protocols, Weber BHF, Langmann T (eds) pp 245–254. Totowa, NJ: Humana Press

Sethna S, Scott PA, Giese APJ, Duncan T, Redmond TM, Riazuddin S, Ahmed ZM (2020) Loss of CIB2 causes non-canonical autophagy deficits and visual impairment. bioRxiv: 2020.09.18.302174

Smith RJ, Berlin CI, Hejtmancik JF, Keats BJ, Kimberling WJ, Lewis RA, Möller CG, Pelias MZ, Tranebjaerg L (1994) Clinical diagnosis of the Usher syndromes. Usher Syndrome Consortium. American journal of medical genetics 50: 32–8

Toms M, Pagarkar W, Moosajee M (2020) Usher syndrome: clinical features, molecular genetics and advancing therapeutics. Ther Adv Ophthalmol 12: 2515841420952194

Travis GH, Golczak M, Moise AR, Palczewski K (2007) Diseases Caused by Defects in the Visual Cycle: Retinoids as Potential Therapeutic Agents. Annual Review of Pharmacology and Toxicology 47: 469–512

Trouillet A, Dubus E, Dégardin J, Estivalet A, Ivkovic I, Godefroy D, García-Ayuso D, Simonutti M, Sahly I, Sahel JA, El-Amraoui A, Petit C, Picaud S (2018) Cone degeneration is triggered by the absence of USH1 proteins but prevented by antioxidant treatments. Scientific Reports 8: 1968

van Roy F (2014) Beyond E-cadherin: roles of other cadherin superfamily members in cancer. Nature Reviews Cancer 14: 121–134

van Wijk E, van der Zwaag B, Peters T, Zimmermann U, Te Brinke H, Kersten FF, Marker T, Aller E, Hoefsloot LH, Cremers CW, Cremers FP, Wolfrum U, Knipper M, Roepman R, Kremer H (2006) The DFNB31 gene product whirlin connects to the Usher protein network in the cochlea and retina by direct association with USH2A and VLGR1. Hum Mol Genet 15: 751–65

Vernon M (1969) Usher’s syndrome--deafness and progressive blindness. Clinical cases, prevention, theory and literature survey. J Chronic Dis 22: 133–51

Wald G, Brown PK (1956) Synthesis and Bleaching of Rhodopsin. Nature 177: 174–176

Weston MD, Kelley PM, Overbeck LD, Wagenaar M, Orten DJ, Hasson T, Chen ZY, Corey D, Mooseker M, Sumegi J, Cremers C, Moller C, Jacobson SG, Gorin MB, Kimberling WJ (1996) Myosin VIIA mutation screening in 189 Usher syndrome type 1 patients. American journal of human genetics 59: 1074–1083

Williams DS, Aleman TS, Lillo C, Lopes VS, Hughes LC, Stone EM, Jacobson SG (2009) Harmonin in the murine retina and the retinal phenotypes of Ush1c-mutant mice and human USH1C. Investigative ophthalmology & visual science 50: 3881–3889

